# Genetic evidence for functions of Chloroplast CA in *Pyropia yezoensis*: decreased CCM but increased starch accumulation

**DOI:** 10.1101/2023.05.22.541789

**Authors:** Baoyu Zhang, Xueying Liu, Xiujun Xie, Li Huan, Hong Wang, Zhizhuo Shao, Guangce Wang

## Abstract

To adapt to the change of intertidal environment, intertidal macroalgae have evolved complicated Ci utilization mechanism. However, our knowledge regarding the CO2 concentrating mechanism (CCM) of macroalgae is limited. Carbonic anhydrase (CA), a key component of CCM, plays important roles in many physiological reactions in various organisms. While there are a large number of genes encoding CA in the Pyropia yezoensis genome, the exact function of specific CA in P. yezoensis remains elusive. To explore the specific function of chloroplast CA in intertidal macroalgae, we produced chloroplast-localized βCA1 knockdown mutants of P. yezoensis through RNA interference, and Pyca1i mutants showed a notable decrease in leaf area and overall biomass, as well as decreased soluble protein and unsaturated fatty acid content under different DIC conditions. However, Pyca1i mutants showed relatively higher starch content compared to the wild-type. The activity of enzymes involved in Calvin cycle, photorespiration, Pentose-phosphate pathway and floridean starch synthesis of P.yezoensis indicated an effective starch accumulation pathway after interference of βCA1. All results suggest that the decreased activity of PyβCA1 impaired the CCM and development of thalli of P.yezoensis, but stimulated starch accumulation in the cytoplasm through feedback to the photorespiration pathway and PP pathway to replenish intermediates for the Calvin cycle. This study is the first to explore the specific function of chloroplast CA in intertidal macroalgae using genomic technology. The results provide valuable insights into the adaption mechanisms of intertidal macroalgae to their environment.

## Introduction

On earth, photoautotrophs utilize the photosynthetic Calvin-Benson cycle to transform inorganic carbon into organic carbon. The crucial enzyme responsible for this conversion in the Calvin cycle is Ribulose-1,5-bisphosphate carboxylase/oxygenase (Rubisco), which employs CO_2_ as a substrate. C4 higher plants employ an efficient carbon sequestration method whereby CO_2_ is initially converted into HCO_3_^-^ by carbonic anhydrase (CA), which then serves as substrate for Phosphoenolpyruvate carboxylase (PEPC). CA is essential for C4 plants (Von Caemmerer et al. 2004). In seawater, the CO_2_ concentration is roughly 2000 times lower than in air, and diffusion is 8,000-10,000 times slower. As a result, HCO_3_^-^ is the main inorganic carbonate in seawater (Zeebe 2011, Young et al., 2012), and marine algae (microalgae or macroalgae), utilize it as carbonic source for growth. Therefore, studying CA is of great significance in comprehending the various forms of inorganic carbon fixation present on Earth.

Most intertidal macroalgae are economically valuable. *Pyropia yezoensis*, a representative species found in the upper intertidal zone, is an important red alga and widely cultivated in East Asian countries such as China, Japan and South Korea, with an annual production of about 1.8 million tons in 2017 (FAO, 2019). The wild leafy thalli of *P. yezoensis* undergo the emersion and submersion with the changing tides every day. Physiological researches indicated that *Pyropia* could utilize atmospheric CO_2_ when emersion, while fix HCO_3_^-^ in seawater when submerged (Zou and Gao 2002, Zhou et al. 2014, Huan et al. 2018). Thus, farmers applied semi-floating raft method and regularly exposed *P.yezoensis* to the atmosphere during marine aquaculture, which improves the quality and yield of *P.yezoensis* (Zhou et al. 2014). Since the thalli of *P.yezoensis* are arranged in monolayer cells (Wang et al. 2004), and are surrounded by varying Ci types, achieving quick conversion between HCO_3_^-^ and CO_2_ in thalli cells of *P.yezoensis* will depend on CAs, thus, CA is essential for *P.yezoensis*.

Diverse subtypes and distributions of CA have been found in plants and algae (Moroney et al. 2001, Ignatova et al. 2019), including *Pyropia*. To cope with the complicated intertidal environment, *Pyropia* has expanded CA genes and anti-oxidative related gene. Wang et al. (2020) suggested that 24 putative CA genes might exist in *P. yezoensis* genome, while 17 CA genes were found in *P. haitanensis* (Chen et al. 2022). Our previous studies have identified 11 CA genes with complete coding sequence, belonging to α-, β- and γ-types. The change of these CA genes with different [HCO_3_^-^] at the transcript level, and the localization of seven CAs has been detected and defined in *P. yezoensis* (Zhang et al. 2020, 2022). However, our previous experiments were based on the cellular level. To deeply explore the functions of specific CA, genetically transformed algal strains are necessary.

Stable genomic transformation systems have been achieved in *P.yezoensis*, and stable mutants have been obtained based on functional genes such as γ-CA1 like (γCAL1), high–light inducible protein (HLIP), one-helix proteins (OHPs) et al. (Zheng et al. 2021, Shao et al. 2022). It has been proved that genomic transformation is a powerful tool for uncovering the functions of target genes. In this study, we focused on chloroplastic βCA1 in the leafy thalli of *P. yezoensis*, and constructed RNA interference mutants of βCA1 by inserting reverse sequence. After verifying these mutants through sequencing, we obtained five mutants. By examining changes in RNA and protein levels, we identified three typical strains - *Pyca1i-1*, *Pyca1i-2* and *Pyca1i-6* - for further study under different bicarbonate concentrations: NC (2 mM NaHCO_3_ in seawater) and HC (8 mM NaHCO_3_). Of the three mutant lines tested under these conditions, *Pyca1i-1* and *Pyca1i-2* mutants showed notable decreases in leaf area and overall biomass. Decreased βCA1 activity had negative effects on DIC affinity and even affected the metabolism of the main storage. This research shed light on the roles of an abundant chloroplast βCA1 in bicarbonate utilization, growth and metabolism in the intertidal macroalga *Pyropia*, providing novel insights into the physiological mechanisms of this species.

## Results

### Target fragment insertion into *P. yezoensis* genome

After being subjected to hygromycin B stress for 8 weeks, five hygromycin B-resistant strains of *P.yezoensis* were isolated and named *PyβCA1i-1*, *PyβCA1i-2*, *PyβCA1i-4* to *6*. Genomic PCR was conducted to verify the presence of *Pycali* expression cassette in the *P. yezoensis* genome, and sequencing was performed to confirm the presence of the expression plasmid with targeting gene sequence, but not in WT (Fig. 1). To assess the stability of the expression cassettes in the *P. yezoensis* genome across generations, the mutants were re-examined after 6 months of culturing through monospores germination (Supplemental Fig. 1). The results indicated that the target fragment was stably inserted into the *P. yezoensis* genome and could be transferred through reproduction.

**Fig. 1.**
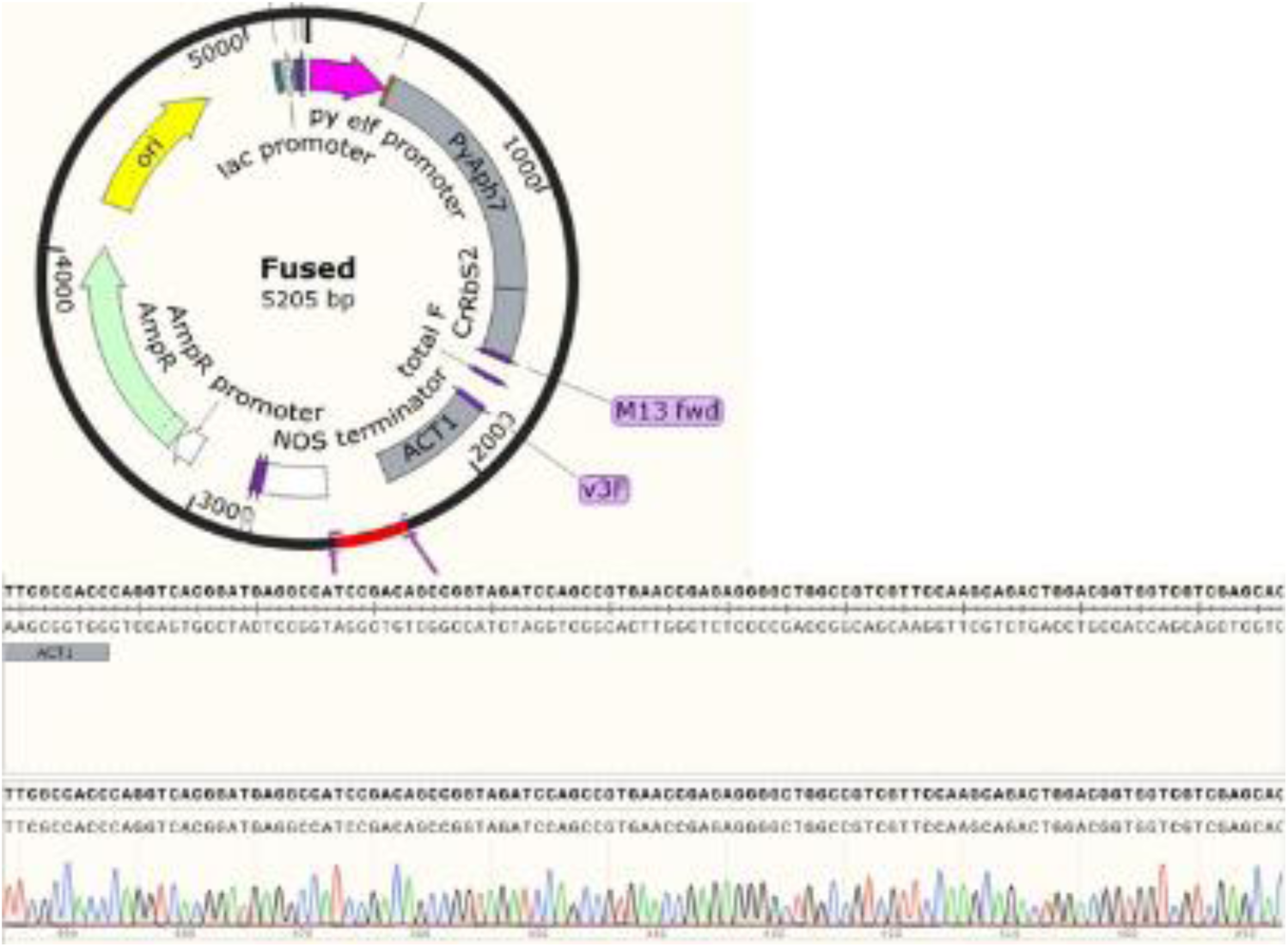
Antisense PyβCA1 targeting site in pEA7 expression plasmid and the sequencing result of *Pyca1i* mutants.

### The change of RNA and protein levels of *PyβCA1i* mutants

The RNA and protein levels of the mutants and WT was detected by RT-qPCR and Western blot, respectively. The RNA level of mutants and WT showed a significant difference, with the *βCA1* gene expression in *Pyβca1i* lines decreasing approximately 5-fold compared to WT (Fig. 2a). An immunoblot analysis of total proteins extracts, using an anti-CA1 monoclonal antibody and a RbcL polyclonal antibody, revealed a positive band at around 28 KD and 54 KD, respectively, which matched the predicted molecular weight of PyβCA1 and RbcL protein. RbcL was used as a standard protein. The protein level of βCA1 were lower in mutants compared to WT (Fig. 2b). On the other hand, immunoblot results also indicated βCA1 was abundant in the 20 μg loading total protein.

**Fig. 2.**
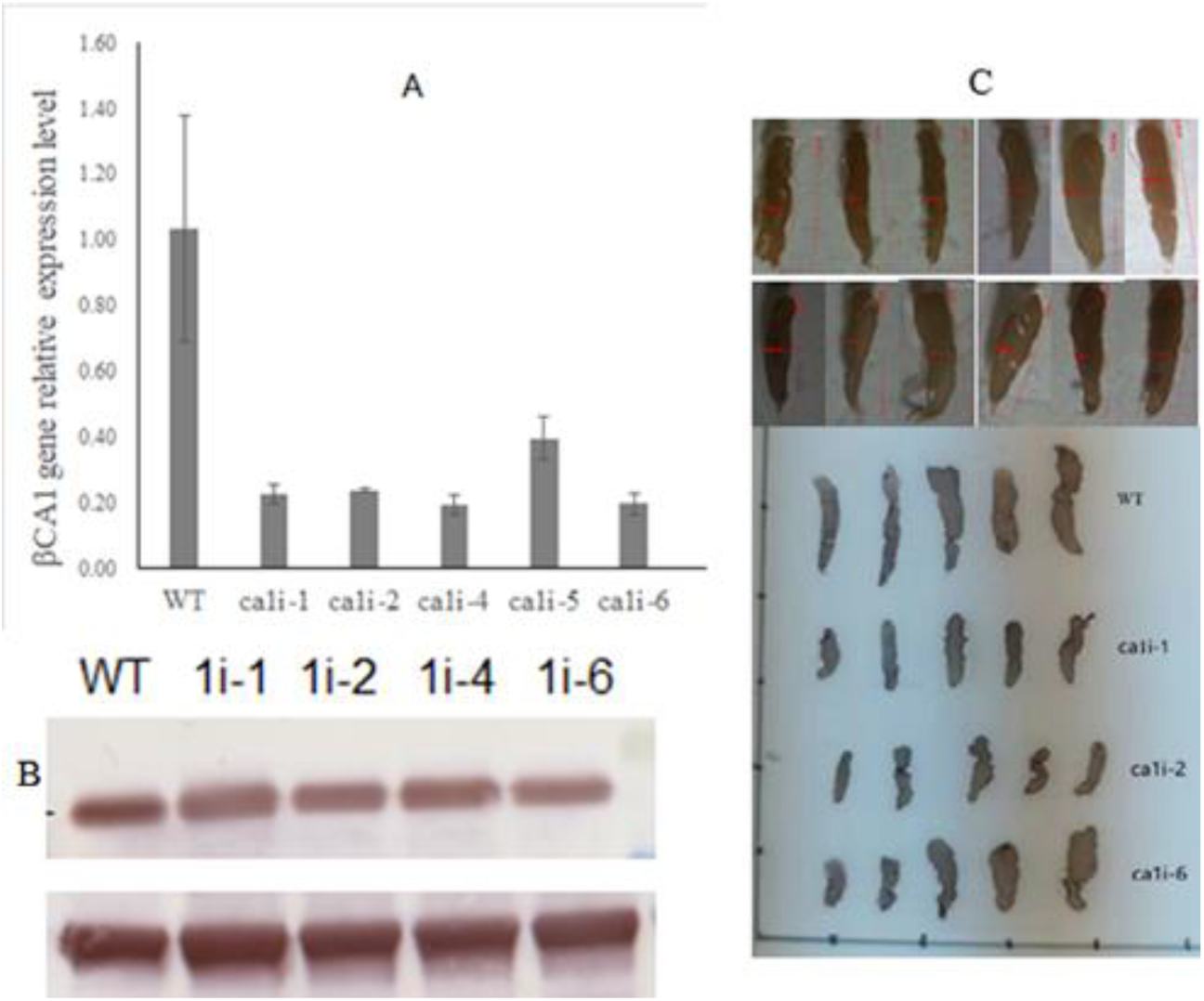
Changes in RNA and protein level of PyβCA1 among Pyca1i mutants and wild type (WT) lines, and phenotypes of *Pyca1i* mutants and WT under the same culture conditions. (A) Change in RNA level of *PyβCA1* between βCA1-antisense mutants and WT. (B) Western blot analysis of PyβCA1 in WT and *PyβCA1i* mutants. Rubisco large-subunit (RbcL) was used as the loading control. (C) Phenotype of *PyβCA1i* mutants and WT after cultivation 20 days under the same culture conditions.

### Morphological characteristics and biomass of *PyβCA1i* strains

Under NC condition (2mM NaHCO_3_), the general morphological and developmental phenotypes of *ca1i-1*, *ca1i*-*2* and *ca1i*-*6* mutants were observed. The mutants, especially *ca1i-1* and *ca1i-2*, were visibly smaller in length and width compared with WT after 20 days of growth under the same conditions. The phenotype of *ca1i-6* was similar to WT (Fig. 2c). Based on the observed differences in RNA and protein levels, as well as morphological characteristics of the mutants, *ca1i-1*, *ca1i-2* and *ca1i-6* were selected for further investigation.

### Changes in pH, photosynthetic oxygen evolution (POE) and respiration rate under different DIC conditions

To investigate the physiological responses of the *βca1i* mutants to HC (8 mM NaHCO_3_), these three strains were cultured under NC and HC conditions, with WT as control.

The pH changes were similar between the *ca1i* mutants and WT under NC conditions, but the highest pH reached during 14 days’ cultivation was different. The pH increased from 8.2±0.02 to 8.9±0.02 for the *ca1i* mutants, and to 9.1±0.02 for WT. However, the pH changes in HC medium were notably faster than in NC, increasing to 9.2±0.02 for *ca1i-1* and *ca1i-2* on the 14^th^ day, whereas it increased to 9.4±0.02 on the 12^th^ day and then decreased to 9.2±0.02 for WT on the 14^th^ day, and the change of pH in *ca1i-6* was similar to WT (Fig. 3a).

**Fig. 3.**
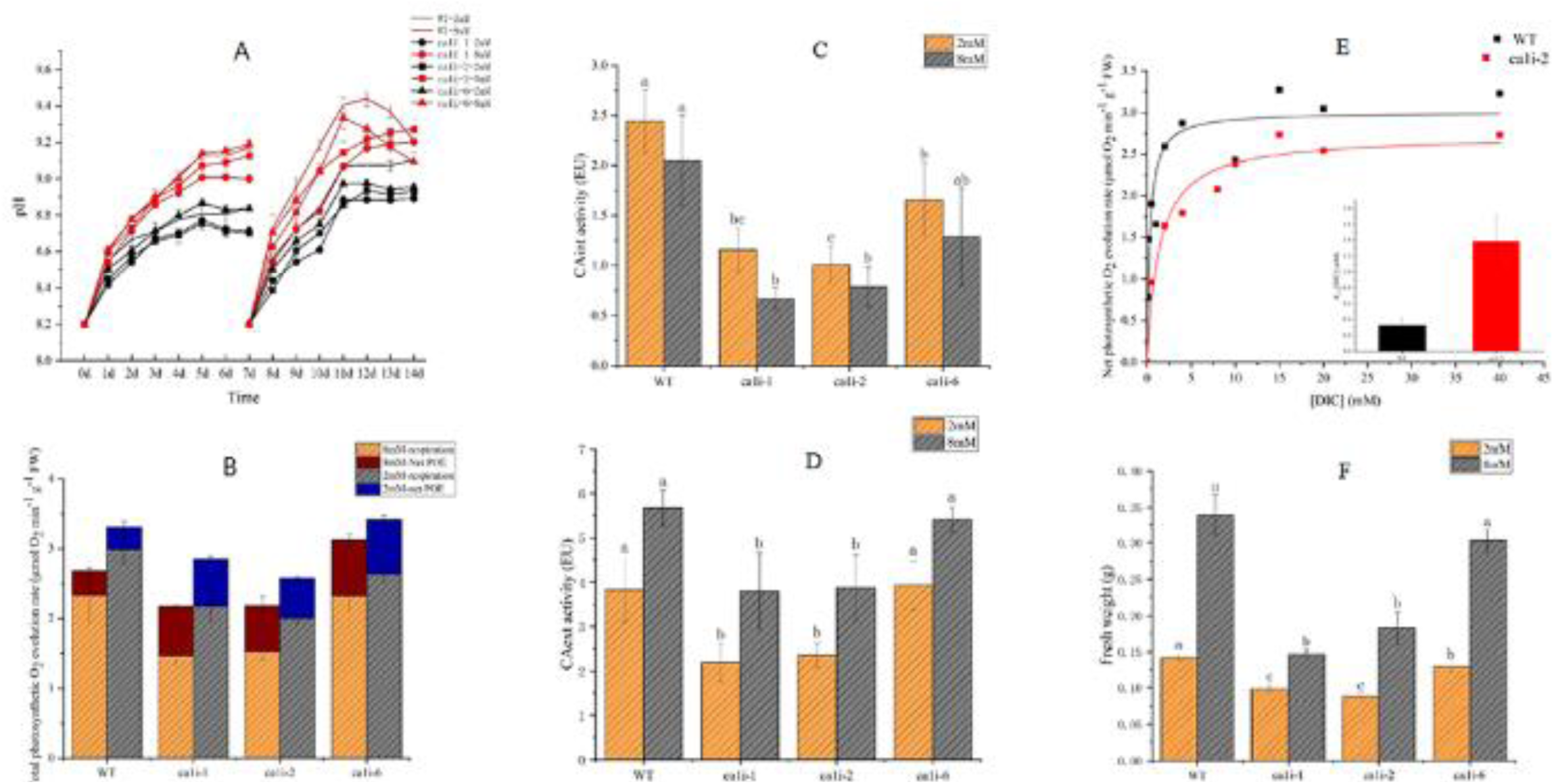
Physiological features and CA activity of *PyβCA1i* mutants and WT under different DIC conditions. (A) pH change in media during culturing mutants and WT, respectively. (B) Photosynthetic oxygen evolution rate and respiratory rate of *Pyca1i* mutants and WT. (C-D) Intracellular and extracellular CA activity in mutants and WT, respectively. Data represent the mean ± standard deviation from three biological replicates. Means followed by the same lowercase letters are not significantly different at p ≤ 0.05 by one-way ANOVA and Tukey’s test. (E) Thalli of *P.yezoensis* photosynthetic DIC affinity assay and calculated [DIC] for half maximal rate = *K_0.5_* [DIC]. (F) Fresh weight of thalli of *P. yezoensis* mutants and WT under different DIC conditions after 14 days cultivation.

The POE rate of mutants was lower than that of WT under both DIC conditions. At pH 8.2±0.02, The POE rate were 1.4 and 2.2 μmol ml^-1^·gFW^-1^·min^-1^ for mutants and WT, respectively, under NC conditions. Under HC conditions, the POE rate increased to 2.0 and 3.0 μmol ml^-1^·gFW^-1^·min ^-1^ for mutants and WT, respectively, (Fig. 3b).

The respiration rate of these three mutant lines were all higher than in WT under both NC and HC conditions, with rate of 0.6 μmol ml^-1^·gFW^-1^·min ^-1^for mutant lines and 0.34 for WT (Fig. 3b).

### Decreased intracellular carbonic anhydrase activity (CAint) and extracellular CA activity (CAext) compared to WT under NC and HC conditions

The basic function of CA is to catalyze the reversible interconversion of CO_2_ and HCO_3_^-^. Compared to WT, the CAint and CAext were decreased in *ca1i* mutants when cultured under NC and HC conditions (Fig. 3c). Specifically, the CAint of WT was 2.1- and 2.4-fold higher than that of *ca1i-1* and *ca1i-2*, respectively, while being 1.5-fold higher than that of *ca1i-6*. However, with increasing of DIC concentration, the CAint of both mutants and WT decreased, with mutants showing a reduction of up to 40%, compared to NC condition. While WT decreased only about 16% compared to NC conditions.

Under HC conditions, the CAext of both mutants and WT increased, with the CAext of WT increased by 48%, and the CAext of *ca1i-1* and *ca1i-2* increasing by approximately 70%, compared to NC condition. However, the CAext of mutants remained lower than that of WT, and the CAext of WT was 1.7-1.5 fold higher than that of *ca1i-1* and *ca1i-2* under NC and HC condition, respectively (Fig. 3d). The CAext and CAint activity of transgenic line *ca1i-6* was similar to that of WT, with no apparent differences between them under the two DIC conditions (Fig. 3d).

### Interference of chloroplast βCA1 impaired Ci affinity and resulted in relative higher residual [**HCO_3_^-^**] in the mutant medium

The half-saturation constant for DIC (*K_1/2_*[DIC]) represents the subject’s DIC affinity, with a higher value indicating lower affinity for DIC. The *K_1/2_*[DIC] in artificial seawater-grown *βCA1i* lines was approximately 4.27-fold higher than that in WT lines (Fig.3e).

On the 7^th^ day, the bicarbonate concentration in medium varied differently between mutants and WT (Table 1). The residual bicarbonate concentration in mutant medium was higher than in WT medium, especially under HC conditions, where the divergence was more evident. After 6 days of cultivation, there was approximately 3mM HCO_3_^-^ in mutant medium, while it was about 1.4 mM HCO_3_^-^ for WT.

**Table 1.**
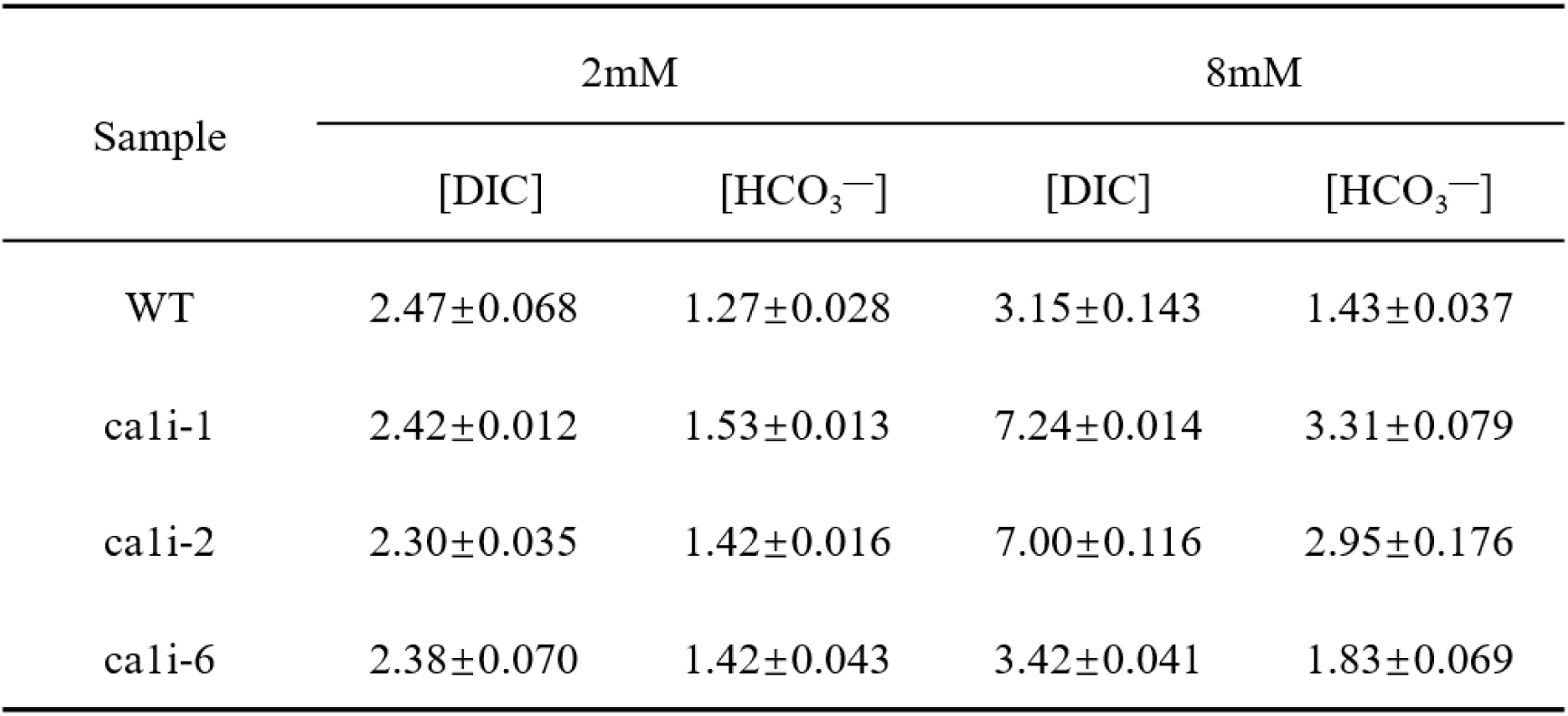
The concentration of DIC and bicarbonate in medium before and after cultivation

### Biomass of *Pyca1i* mutants cultured under NC and HC conditions

Regarding biomass, the growth of *ca1i-1* and *ca1i-2* mutants was retarded relative to WT, resulting in a 31-38% lower biomass than WT under NC, while *cali-6* lowered biomass approximately 9%. Under HC conditions, *ca1i-1* and *ca1i-2* mutants showed a 46-56% lower biomass than WT, while *cali-6* lowered biomass by only 11% (Fig. 3f).

### The total soluble protein, fatty acid (FA) and starch content in mutants and WT under different DIC

*P. yezoensis* is a macroalga widely known for its high nutritional value, particularly its rich in protein, amino acids and unsaturated fatty acid (FA). We compared the total soluble protein, FA and starch content of *Pyca1i* mutants and WT under different DIC conditions after 14 days of cultivation (Fig. 4).

**Fig. 4.**
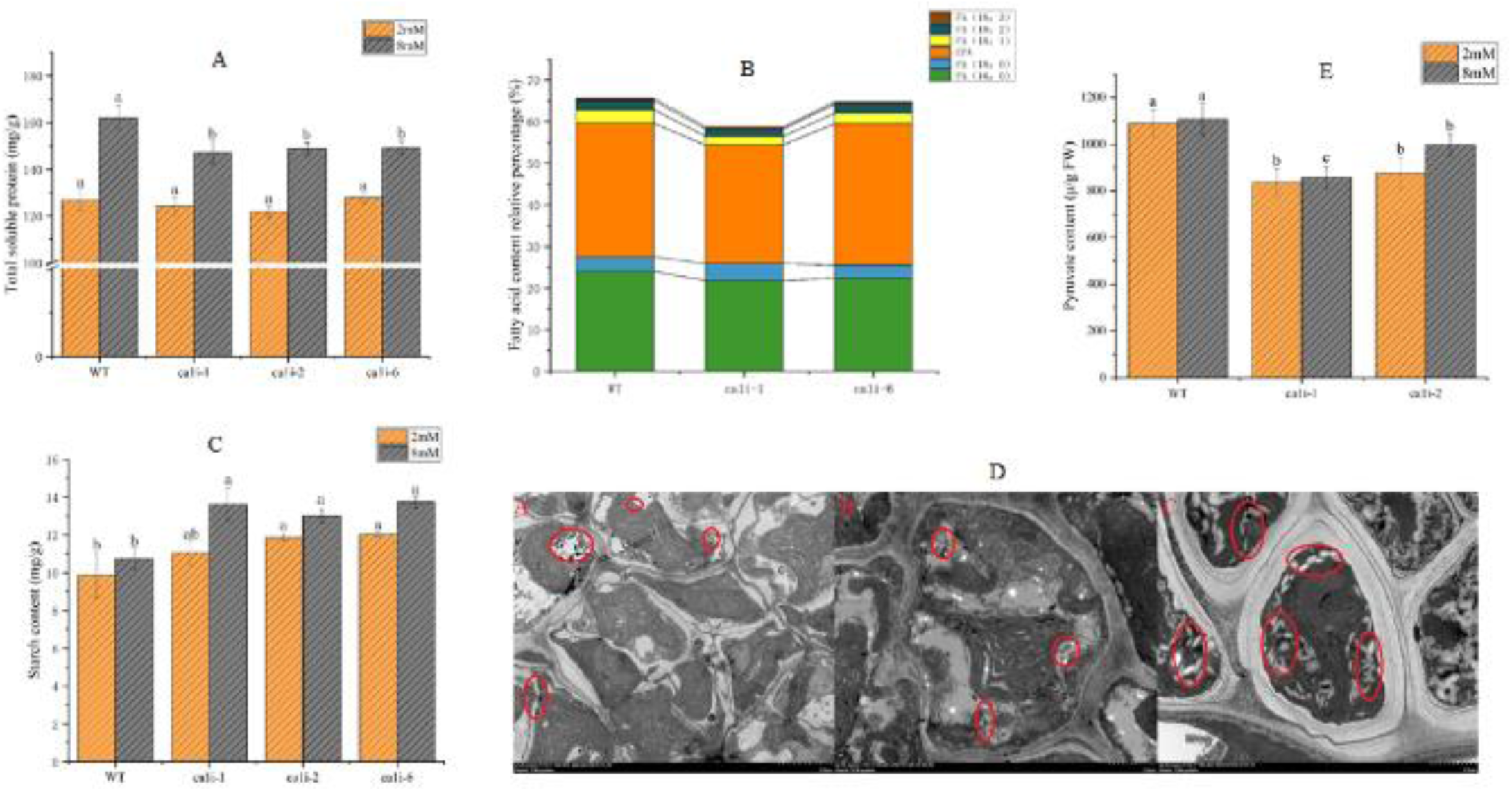
Changes in main metabolites content in mutants and WT. (A) Total soluble protein content among mutants and WT under different DIC conditions. (B) Relative content of fatty acid components in mutants and WT. (C) Starch content of mutants and WT under different DIC conditions. (D) Transmission electron microscopy images of mutants and WT thalli under normal cultivation conditions. (E) Pyruvate content of mutants and WT under different DIC conditions. Data represent the mean ± standard deviation from three biological replicates.

Under NC conditions, the total soluble protein was similar for both mutants and WT, at around 125 mg/g DW. However, under HC conditions, the total soluble protein content increased for mutants to 146 mg/g DW, while WT still had a higher content of 161.97 mg/g DW (Fig. 4a).

We also analyzed the FA composition by gas chromatography and found that mutants had lower levels of C16:0, the main saturated fatty acid in *Pyropia*, compared to WT, with levels of 21.8-22.3%, and 24.1%, respectively. Additionally, mutants had lower levels of EPA, the main unsaturated fatty acid, than WT at 28.4-30% and 32.1%, respectively, under NC conditions (Fig. 4b).

The starch content for mutants and WT was 11.06-12.02 mg/g DW and 9.85 mg/g DW, respectively, under NC conditions. However, under HC conditions, the starch content increased to 12.98-13.73 mg/g DW for mutants and 10.73 mg/g DW for WT (Fig. 4c). This was further confirmed by transmission electron microscopy (TEM) images of starch accumulation in both *Pyca1i* mutants and WT under NC conditions (Fig. 4d).

### Pyruvate content in mutants and WT under NC and HC conditions

Under the two DIC conditions, pyruvate content was significantly lower in mutants compared to WT (P< 0.05), and did not increase much with higher [HCO_3_^-^] concentration in the medium (Fig. 4e).

### Activity of enzymes involve in floridean starch synthesis, PP pathway and photorespiration showed total divergence between *ca1i* mutants and WT

To account for changes in pH under two DIC conditions, with pH levels not exceeding 9.0 ±0.1 on the 3^rd^ day, all enzymes activities were detected during cultivation on the 3^rd^ day.

The Rubisco carboxylation activity was similar for both mutants and WT under NC conditions, but increased with higher [HCO_3_^-^] concentration in WT while showed a slight decrease in mutants (Fig. 5a). Glyceraldehyde-3-phosphate dehydrogenase (GAPDH) is a ubiquitous enzyme in plants, with two subtypes, one participates in glycolysis in cytosol and the other involves in the Calvin cycle in plastids. In chloroplast, GAPDH belongs to NADPH-dependent subtype (Zaffagnini et al. 2013). NADPH-GAPDH catalyzes 1,3-Bisphosphoglycerate into 3-phosphoglyceraldehyde in the Calvin cycle and plays a central role in CO_2_ utilization. The activity of NADPH-GAPDH in mutants was significantly higher than in WT under both DIC conditions, with activity ranging from 5.7-6.5 nmol/min/mg prot to 8.0-10.3 nmol/min/mg prot with HCO_3_^-^ increase. In contrast, its activity in WT raised from 3.3 to 4.5 nmol/min/mg prot from NC to HC conditions. The activity of NADPH-GAPDH in mutants was about one fold higher than in WT (Fig. 5b).

**Fig. 5.**
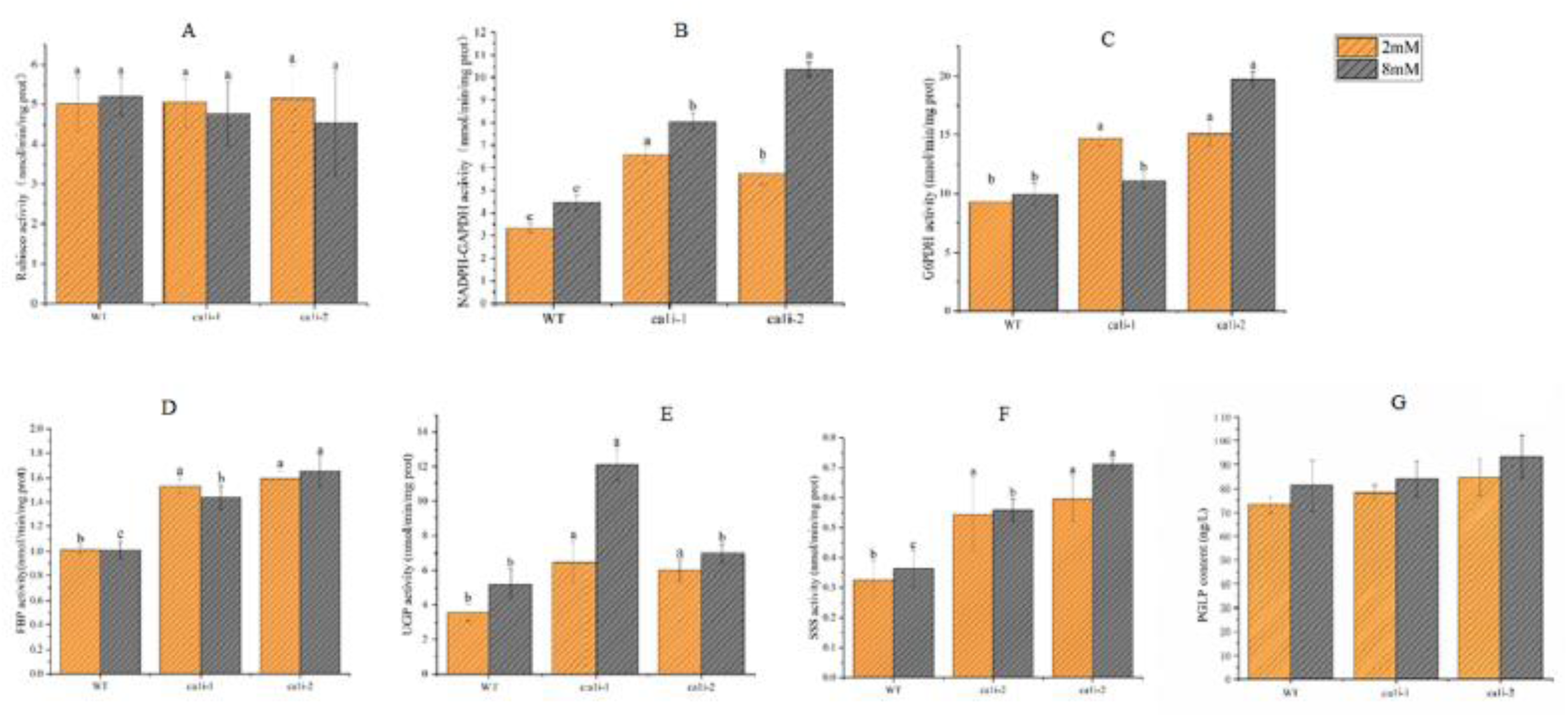
Activities of enzymes involved in Pentose-phosphate pathway, Calvin cycle, photorespiration and floridean starch synthesis. Data represent the mean ± standard deviation from three biological replicates. (A) Rubisco carboxylase activity. (B) NADPH-GAPDH activity. (C) G6PDH activity. (D) Fructose 1,6 bisphosphatase (FBP) activity. (E) UDPG pyrophosphorylase (UGP) activity. (F) Starch synthase (SSS) activity. (G) 2-Phosphoglycolate phosphatase (PGLP) content.

The activities of Glucose-6-phosphate dehydrogenase (G6PDH), which is involved in Pentose-phosphate (PP) pathway were significantly higher in *Pyca1i* mutants than WT under two DIC conditions, with activity approximately 15 nmol/min/mg prot and 9.28 nmol/min/mg prot in mutants and WT, respectively, under NC conditions. With the increase of [HCO_3_^-^] in medium, the enzyme activity increased as well (Fig.5c).

Fructose-1, 6-bisphosphatase (FBP) catalyzes fructose-1, 6-bisphosphate conversion into fructose-6-P (Fru-6-P). The FBP activity in mutants and WT were approximately 1.5-1.6 and 1.0 nmol/min/mg prot, respectively, under NC condition, and this enzyme activity in mutants and WT under HC condition was similar to that under NC (Fig. 5d).

UDPG pyrophosphorylase (UGP) and starch synthase (SSS) participate in the formation of carbonate storage, such as starch and floridoside, of *Pyropia* (Yu et al., 2021). The UGP enzymes activity in mutants and WT was approximately 6 and 3.55 nmol/min/mg prot under NC condition, respectively, and increased to approximately 10 and 5 nmol/min/mg prot under HC condition, for mutants and WT, respectively (Fig. 5e). The change in SSS activity was similar to that of UGP activity between mutants and WT, furthermore, the SSS activity in *Pyca1i-2* was about 1-fold higher than that of WT under these two DIC conditions (Fig. 5f).

2-Phosphoglycolate phosphatase (PGP) catalyzes the initial step of the photorespiratory pathway in various organisms, including yeast. The PGP content under NC conditions was measured to be 73.19 ng L^-1^ for WT and around 80 ng L^-1^ for the mutants. However, under HC conditions, the PGP content in the mutants increased to approximately 90 ng L^-1^ when the [HCO3-] in the medium was increased, while the PGP content in WT remained lower than that of the mutants, measuring 81.2 ng L^-1^ (Fig. 5g).

## Discussion

### Chloroplast-localized abundant CAs plays a more important role in photosynthesis and growth of aquatic photosynthetic species than in terrestrial C3 plants

This study showed that the increase in CAext activity in wild type corresponded to the increase in HCO_3_^-^ concentration in the medium (Fig. 3d), indicating that *P.yezoensis* employs biophysical CCM by utilizing bicarbonate in seawater, consistent with previous reports (Zou and Gao 2002, Moulin et al. 2011, Zhang et al. 2020, 2022). Inorganic carbon concentration in seawater increased, as a result, leading to a decrease in CAint activity of *P.yezoensis* wild type (Fig. 3c), which is a characteristic of CCM. CCM is induced by ambient CO_2_ in aquatic environment, but it is not active under higher CO_2_ (i.e., 30,000-50,000 ppm CO_2_) (Wang et al. 2015, Mackinder et al. 2017, Wei et al. 2019).

As an abundant chloroplast-localized CA (Zhang et al. 2020, Fig. 2b), decreasing CAint activity by 40-50% in *Pyca1i* lines compared with WT (Fig. 3c) led to an obvious decrease in POE rate (Fig. 3b), lower affinity for bicarbonate (Fig. 2e), and even retarded growth (Fig. 2c and 3f). Similar results were also observed in some microalgae such as *C. reinhardtii*, *P. tricornutum* and *Nannochloropsis oceanica* (Wei et al. 2017, Gee and Niyogi 2017). The decrease in the thylakoid lumenal CA (CAH3) activity in *C.reinhardtii* severely impaired photosynthesis and normal physiological development, particularly under ambient CO_2_ (Karlsson et al. 1998, Markelova et al. 2009). Another abundant β-CA in *C. reinhardtii*, originally identified as low CO_2_ - inducible B (LCIB), was dispersed throughout the stroma under ambient CO_2_ (Yamano et al. 2021), and played critical roles in the HCO_3_^-^ uptake; its deletion also affected normal growth under ambient CO_2_ (Adler et al. 2022).

In contrast to microalgae and macroalgae, C3 plants require minimal CA activity (Badger and Price, 1994). For instance, deleting two stromal CAs in tobacco showed no effect on photosynthesis (Hines et al. 2021). This phenomenon was also observed in *Zea mays* double CAs antisense mutants (Crawford and Cousins 2022) and *Arabidopsis* deficient α-CA2 mutants (Zhurikova et al. 2016). All of these studies indicate that the activity of chloroplast-localized CAs, especially those that are highly abundant, play more crucial roles in the photosynthesis and growth of aquatic algae than in that of terrestial C3 plants.

### The interference of βCA1 impaired the CCM, which decreased the assimilation of bicarbonate and reduced the content of total soluble protein and FA, but tiggered a feedback mechanism that led to the accumulation of floridean starch

The relative high residual bicarbonate concentration in *ca1i* mutants medium under HC conditions (Table 1), and the increased *K_1/2_*[DIC] (Fig. 2e) indicate that the lower intracellular bicarbonate concentration in mutant lines, particularly under HC conditions, may have affected their metabolic pathways. Besides being catalyed by CA and turns into CO_2_, HCO_3_^-^ also serves as substrates in various pathways, such as amino acid metabolism, fatty acid synthesis and elongation, and leucine catabolism (Bauwe 1986, Nikolau et al. 2003). Pyruvate is a key intermediate in several metabolic pathways, such as gluconeogenesis and TCA cycle, and is essential for the synthesis of FA and protein (Nikolau et al. 2003). The synthesis of malonyl CoA and long-chain fatty acid primarily occurs in plastids (Raven 1995, Brawley et al. 2017). The relative lower pyruvate content in mutants (Fig.4e) led to the lower FA and total soluble protein in mutants (Fig.4a, 4b). Although high crude protein and unsaturated FA content in *Porphyra* (later renamed as *Pyropia*) are one of reasons for its popular (Fleurence1999, Noda 1993, Blouin et al. 2006).

Rubisco is a key component of Calvin cycle and can play carboxylation or oxygenation, depending on the ratio of CO_2_:O_2_ around it, The intermediates of Rubisco carboxylation, triose phosphate, including G3P and DHAP, are substrates for starch synthesis in plants and algae. Floridean starch, floridoside, and mannitol are the main storage carbohydrates of most red algae (Rioux et al. 2015, Martinez-Garcia M & van der Maarel 2016); furthermore, the red algae produce granulated floridean starch in the cytoplasm, which is different from green algae and higher plants that store starch in chloroplast (Viola et al. 2001, Bates et al. 2013). Fünfgeld and colleagues recently proved that ADPG pyrophosphorylase (AGP) in the chloroplasts of *Arabidopsis* was needed for starch synthesis in plastid, but not SSS in the cytosol (Fünfgeld et al. 2022). In contrast to higher plants, it is UDPG pyrophosphorylase (UGP), SSS, branching enzyme (BE) and isoamylase that contribute to the formation of carbonates storages in green and red algae (Patron and Keeling 2005, Yu et al. 2021).

Although there is no significant difference in Rubisco carboxylation between WT and mutants under two DIC conditions, mutants showed a slightly lower Rubisco carboxylation activity under HC conditions (Fig.5a). However, the starch content in mutants was higher than in WT under NC and HC conditions (Fig.4c). This suggests that there may be another pathway for starch accumulation or other ways to replenish the intermidates of carbohydrate storage formation. The higher NADPH-GAPDH activity in mutants (Fig. 5b) indicates that there is ample substrate, 3-PGA, for its reaction in the Calvin cycle. In addition to the pathway from Rubisco carboxylation, photorespiration from Rubisco oxygenation may also replenish 3-PGA, which was coincided with relatively higher 2-Phosphoglycolate phosphatase (PGLP) content in mutants (Fig. 5g). PGLP is a key enzyme in photorespiration and catalyzes the first reaction photorespiration C2 cycle. Flügel et al. (2017) proved that t he change of PGLP activity in photorespiration affects the content of 2-phosphoglycolate, and then affects photosynthesis and starch accumulation in *Arabidopsis*. PGLP1 overexpression lines showed significantly starch content than WT; while lower starch content in antisense plants when they expose to normal or low CO_2_. PGLP is also indispensible in C4 plants to maintain carbon assimilation and allocation (Levey et al. 2018)

Triose phosphate can enter cytoplasm through chloroplast membranes and serve as a substrate for starch and carbonate storage synthesis (Viola et al.2001). The enzymes FBP, UGP and SSS play key roles in this process, which results in the production of starch and floridoside. In the mutants, the higher activitiy of these enzymes (as shown in Fig. 5d, 5e, 5f) supports the higher starch content compared to WT. However, the outflow of triose phosphate from the chloroplast could lead to a decrease in the supply of RuBP, a key substrate for Rubisco. The higher activity of G6PDH in mutants (Fig. 5c) suggests that it replenishes RuBP.

In summary, the decrease of βCA1 activity impaired CCM, development, and total soluble protein and fatty acid content, but stimulated starch accumulation in the cytoplasm by feeding back such pathways as photorespiration and PP pathway to replenish intermediates for the Calvin cycle (Fig. 6). Although the mutants lines had a relatively lower [HCO_3_^-^], the higher catalytic activity of CA did not result in an obvious decrease of Rubisco carboxylation.

**Fig. 6.**
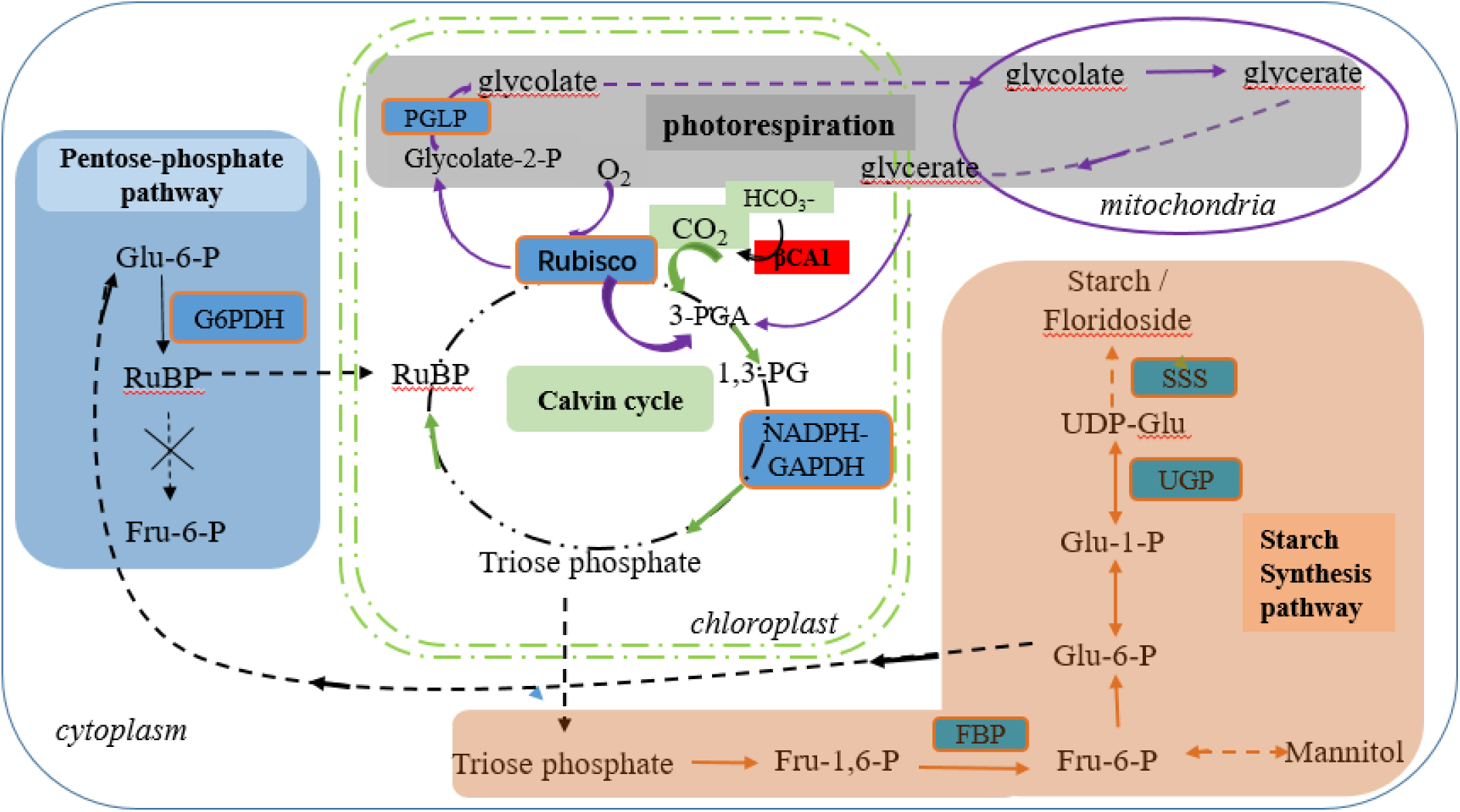
Schematic diagram of pathways which contribute to floridean starch synthesis in *P.yezoensis*, after interference of chloroplast-localized βCA1.

While our previous work found a large number of CA isoforms in *Pyropia* (Zhang et al. 2020, 2022), Wang et al. (2020) predicted that five α-CAs and two βCAs were targeted to the plastids. However, given the potential inaccuracy of predictions, it is possible that there are multiple CAs in the plastids. Our findings indicate that the activity of other chloroplast CAs in *P. yezoensis* cannot fully compensate for the decreased activity of βCA1 during development. In addition, intertidal seaweed may employ other pathways to cope with the decrease in chloroplast-localized CA.

## Materials and Methods

### Algal materials and different DIC setting

Wild type leafy thalli of Pyropia yezoensis was generated from sporophytes of this species, stored in the algae collection in the Institute of Oceanology, Chinese Academy of Sciences, Qingdao, China. The thalli were cultured in Provasoli’s enriched seawater medium at 16±1°C under 12 -h photoperiod at 30 μmol·m^2^·s ^-1^.

To test the reaction of leafy thalli to different [DIC], artificial seawater with different concentration of NaHCO_3_ was used, where 2mM and 8 mM NaHCO_3_ represented NC and HC condition, respectively.

### Expression plasmid construction and algal transformation

Genetic transformation of P. yezoensis was carried out as previously described (Zheng et al. 2021) with slight modifications. The expression plasmid pEA7-PyAct1::PyCFP was kindly provided by Prof. Toshike Uji from Kokkaido University (Hirata et al. 2014). To create the *PyβCA1* RNAi expression cassette, *PyCFP* gene in this plasmid was replaced with 220 bp reverse complementary coding region of *PyβCA1* gene. This region was corresponding to partial putative active domain and amplified with the following primers, P7-βCA1i F: 5’-gggaaattcgagctcGCTGACGCACATCCGTGATG-3’, P7-βCA1i R: 5’-gtcaccttcgccaccCAGGTCACGGATGAGGCCAT-3’. The target sequence and vector via In-Fusion cloning technique, and the plasmid pEA7-PyAct1::PyβCA1i was finally produced according to the schematic diagram in Figure1. Particle reparation and bombardment were performed according to previous reported (Zheng et al. 2021, Shao et al. 2022). For each shooting, 10μL of the suspension was used, and the protocol of particle bombardment was the same as that described in Zheng et al. (2021).

### Genomic PCR and qRT-PCR

After 8 weeks selection with hygromycin B, a single thallus was chosen and cultivated in each cell culture flask. Genomic DNA was extracted from each cell culture flask with Plant genomic DNA kits (Tiangen BioTech., Beijing, China). Genomic PCR (gPCR) was conducted with the primer pair F: 5’-CGCATAAGCTTCCGCTCGA-3’ and R: 5’-CTGAGAGTGCACCATAAATTCCC-3’ to test the insertion of the fragment (including partial PyAct1 promoter sequence, PyβCA1i interference region, and partial the NOS terminator sequence) into the genome of P. yezoensis.

To examine the expression level of the transformants, quantitative real-time PCR (qRT-PCR) was performed with PyGAPDH as the interference genes according to Wu et al. (2013). Total RNA was extracted using the Plant Total RNA extraction Kits (Tiangen BioTech) from both WT and mutants, and 1 μg total RNA was used to inverse transcribe to cDNA (Promega Biotech, Madison, WI, USA), which was subsequently used as templates. The protocol for qRT-PCR was similar to that described in Zhang et al. (2020), and the primer pairs for qRT-PCR were as follows: F: 5’-CCA GTC TGC TTG GAACGACG-3’, R: 5’-TAGACCGAATCGATGCCCTC-3’.

### Immunodetection of PyβCA1

Western blotting was performed following the protocol described in Zhang et al. (2022). Specifically, 20 μg of total soluble protein from WT and mutants were loaded per lane and separated in 12% SDS-PAGE. Monoclonal primary antibody against PyβCA1 was prepared by Youke Biotech. Co. (Shanghai, China), while ployclonal antibody against RbcL for plants and algae were obtained from Agrisera (Product No. AS03037). Anti-PyβCA1 or anti-RbcL secondary antibodies were either anti-mouse IgG, HRP conjugated or anti-rabbit IgG, HRP conjugated (GenStar).

### pH change and fresh weigh under different DIC conditions

The pH was monitored daily using a pH meter (Sartorius, Beijing, China) and the medium was changed weekly. On the 14th day, leafy thalli were harvested through a cell filter (Ф100 μm) and redundant water was removed through filter paper. The samples were then weighed and frozen at -80℃ for further use.

### DIC affinity curves and half-saturation determination

DIC-dependent O_2_ evolution was measured using a Clark-type Oxygen Electrode (Hansatech Instruments, UK) at 50 μmol·m2·s-1 and 15±0.1℃. Leafy thalli were cut into small pieces (ca. 2mm × 2mm) and incubated in artificial seawater (ASW) without NaHCO3 under normal cultivation conditions for 24h to minimize the effect of cutting damage on photosynthesis. Approximately 10 mg of sample was transferred to a chamber containing 2mL DIC-free ASW, and reaction media were magnetically stirred. NaHCO_3_ was injected into the chamber when no further O_2_ evolved, creating various DIC concentration by adding aliquots of NaHCO_3_ stock solution into the media. O_2_ evolution was recorded within 10 min after the addition of NaHCO_3_. Photosynthesis curves were processed using a Python (http://www.python.org/) script that utilized the scipy ‘curve_fit’ function to fit a Michaelis-Menten equation to the data and solve for Vmax and *K_1/2_*[DIC].

### Photosynthetic oxygen evolution rate (POE), respiration rate under different DIC conditions

The net POE and respiration rate of WT and mutants was also determined by a Clark-type oxygen electrode as described above, and respiration rate was measured for a period of 10 min after dark.

### Measurement of total DIC and HCO_3_^-^ concentration

Total alkalinity of ASW was determined by titration of seawater at 32 (PSU) and 15±0.1℃ using the method described by Millero et al. (1993), and total [DIC] in the medium after 6 days of cultivation was measured using LI-7000 CO_2_/H_2_O analyzer (LI-COR, USA). All data, including temperature, alkalinity, pH and total [DIC], were input into CO2SYS (Pierrot et al. 2006) and used to calculate HCO_3_^-^ concentration.

### Carbonic anhydrase activity assay

CA activity was carried out according to the methods of Wang et al. (2019) and Fernández et al. (2018) with slight modifications. For external CA (CAext) activity, the time required for the pH to drop from 8.4 to 7.9 was recorded at 4-5℃ using a chamber containing 5 mL of cold Tris-Cl buffer (50 mM tris-cl, 5mM EDTA, 2mM DTT, pH 8.5±0.01, 4℃). Approximately 0.01g fresh leafy thalli was carefully prepared by removing any excess water from the surface with filter paper. Next, 2mL of cold (4℃) CO2-saturated milliQ water (18.3 MΩ cm) was added to the chamber to initiate the reaction. Enzyme activity was expressed in enzyme unit (EU), calculated using the formula: EU =10× (T0 -T)/T, where T0 and T represent the time in seconds for pH drop without and with tissue, respectively. To determine total CA activity, both internal and external, crude extracts were obtained by grinding 50 mg of the sample with sample buffer used for extracellular activity determination. The resulting 150 ul homogenate was added to 4.85 mL Tris-cl buffer and the total CA activity was measured in the same manner as for CAext. Intracellular CA (CAint) was calculated by subtracting CAext activities from total CA activity.

### Measurement of total soluble protein, starch, pyruvate and PGLP content

The leafy thalli were subjected to freeze-drying and used to measure the total soluble protein and starch content. Approximately 30 mg of dried powder was extracted according to the Plant total protein kit protocol (Solarbio Sci & Tech Co. Ltd, Beijing). Similarly, 30 mg dried powder was extracted according to the protocol of Total starch content Kit (Geruisi-bio. Co. Suzhou, China). and the absorbance was measured at 510 nm. The total starch content was calculated by multiplying the glucose content by the conversion coefficient of 0.9.

The pyruvate content in the leafy thalli of WT and mutants was quantified using the Plant Pyruvic Acid Content Kit (Grace Biotech, China) according to the provided protocol. Pyruvic acid reacts with 2,4-dinitrophenylhydrazine to form 2,4-dinitrophenylhydrazone, which exhibits a brownish-red color in an alkaline solution. The absorbance value at 520 nm was measured to determine the pyruvic acid content. Similarly, the PGLP content in the leafy thalli of WT and mutants was assessed using the PGLP Content ELISA Kit (Shanghai Enzyme-Linked Biotech. Co. Ltd., China) following the instructions provided. The absorbance value at 520 nm was measured for quantification purposes

### Free fatty acid (FA) analysis

The total lipid was extracted from 30 mg freeze-dried leafy thalli using a chloroform-methanol method described by Wu et al. (2015).The crude lipid samples were dried with N2 flow until a constant weight was obtained. The fatty acid compositions of different DIC cultivated mutants and WT leafy thalli were determined by gas chromatography.

### Enzyme assay

The enzymatic activities of Rubisco, NADPH-Glyceraldehyde-3-phosphate dehydrogenase (GAPDH), glucose-6-phosphate dehydrogenase (G6PDH), Fructose-1, 6-bisphosphatase (FBP), starch synthase (SSS), UDPG pyrophosphorylase (UGP) were measured using quantification kits (Grace Biotech, China) following the user’s manual. In the protocol, about 100 mg fresh algal materials were ground on ice, and suspended with extraction buffer. After concentration, supernatants were incubated on ice for further enzyme assays. Enzyme activities were determined using microplate readers (Infinite M1000 pro, Sweden) by measuring the change in absorbance at 340 nm. The total protein concentration of supernatants was measured by absorbance at 562 nm with the BCA protein assay kit (GenStar). The enzyme activity unit was defined as nmol NAD(P)H oxidation or NAD(P)+ reduced per minute per gram fresh weight or total protein. The SSS activity was determined by measuring the change in absorbance at 450 nm according to the kit protocol.

### Statistical analysis

For all measurements of metabolites and enzyme activities, one-way ANOVAs with post-hoc Tukey test analyses were performed using IBM SPSS Statistics 23 (IBM Co., Aemonk, NY, USA). Statistical significance was set at P≤0.05. The graphs were plotted using Origin2018 (OriginLab Co., Northampton, MA, USA).

### Legends for figures, tables and supporting information

Supplemental Fig. 1. Electrophoresis pattern of the partial fragment of *PyβCA1* amplified with the P7-βCA1i F / P7-βCA1i R primers, along with the corresponding sequencing results.

## Authors’ contributions

Zhang BY conceived, designed research, and obtaining the mutants; Liu XY and Shao ZY did the physiological experiments and made Tables. Xie XJ did the statistical analysis of this work. Huan L cultivated the materials and measured the enzymes. Wang GC gave constructie suggestions and guidance, discussed the results and revised the manuscript. All authors read and approved the final manuscript.

## References

Adler L, Díaz -Ramos A, Mao Y, Pukacz KR, Fei C, McCormick AJ. (2022) New horizons for building pyrenoid-based CO_2_-concentrating mechanisms in plants to improve yields. Plant Physiol. 27, 190(3):1609–1627. doi: 10.1093/plphys/kiac373.

Badger MR, Price GD. (1994) The Role of Carbonic anhydrase in photosynthesis. Annu Rev Plant Physiol Plant Mol Biol 45: 369–392

Bauwe H, Chollet R. (1986) Kinetic Properties of Phosphoenolpyruvate Carboxylase from C3, C4, and C3-C4 Intermediate Species of Flaveria (Asteraceae). Plant Physiology, 82(3): 695–699, https://doi.org/10.1104/pp.82.3.695

Bates PD, Jewell JB, & Browse J. (2013) Rapid separation of developing Arabidopsis seeds from siliques for RNA or metabolite analysis. Plant Methods 9:9. https://doi.org/10.1186/1746-4811-9-9

Blouin N, Calder B L., Perkins B. Brauley SH. (2006) Sensory and fatty acid analysis of two Atalantic species of Porphyra (Rhodophyta). J. Applied Phycol 18:79–85

Brawley SH, Blouin NA, Ficko-Blean E, et al. (2017) Insights into the red algae and eukaryotic evolution from the genome of *Porphyra umbilicalis* (Bangiophyceae, Rhodophyta). Proceedings of the National Academy of Sciences 114 (31): E6361∼E6370

Chen H, Chu J, Chen J, Luo Q, Wang H, et al. (2022) Insights into the ancient adaptation to intertidal environments by red algae based on a genomic and multiomics investigation of *Neoporphyra haitanensis*. Mol. Biol. Evol. 39(1): msab315 doi:10.1093/molbev/msab315

Crawford JD, Cousins AB. (2022) Limitation of C4 photosynthesis by low carbonic anhydrase activity increases with temperature but does not influence mesophyll CO_2_ conductance. Journal of Experimental Botany 73(3):927–938, https://doi.org/10.1093/jxb/erab464

FAO. (2019) FAO Yearbook of Fishery and Aquaculture Statistics. http://www.fao.org/fishery/static/Yearbook/YB2017_USBcard/index.htm.

Fleurence J. (1999) Seaweed proteins: biochemical, nutritional aspects and potential uses. Trends in Food Technol 10: 25–28

Flügel F, Timm S, Arrivault S, Florian A, Stitt M, et al. (2017) The photorespiratory metabolite 2-Phosphoglycolate regulates photosynthesis and starch accumulation in *Arabidopsis*. The Plant Cell 29: 2537–2351, www.plantcell.org/cgi/doi/10.1105/tpc.17.00256

Fünfgeld MMFF, Wang W, Ishihara H, Arrivault S, Feil R, Smith AM, Stitt M, et al. (2022) Sucrose synthases are not involved in starch synthesis in *Arabidopsis* leaves. Nature Plants 8: 574–82. https://doi.org/10.1038/s41477-022-01140-y

Gee CW, Niyogi KK (2017) The carbonic anhydrase CAH1 is an essential component of the carbon-concentrating mechanism in *Nannochloropsis oceanica*. Proc Natl Acad Sci USA 114: 4537–4542

Hines KM, Chaudhari V, Edgeworth KN, Owens TG, Hanson MR (2021) Absence of carbonic anhydrase in chloroplasts affects C_3_ plant development but not photosynthesis. Proc Natl Acad Sci USA 118: e2107425118

Huan L, Wang C, Gao S, He L, Lu X, Wang X, Liu X, Wang G (2018) Preliminary comparison of atmospheric CO _2_ enhancement to photosynthesis of *Pyropia yezoensis* (Bangiales, Rhodophyta) leafy thalli and filamentous thalli: Carbon uptake after CO_2_ enhancement. Phycological Res 66: 117–126

Ignatova L, Rudenko N, Zhurikova E, Borisova-Mubaraksh M, Ivanov B. (2019) Carbonic anhydrases in photosynthesizing cells of C_3_ higher plants. Metabolites 9,73; 10.3390/metabo9040073

Karlsson J (1998) A novel alpha -type carbonic anhydrase associated with the thylakoid membrane in *Chlamydomonas reinhardtii* is required for growth at ambient CO_2_. The EMBO Journal 17: 1208–1216

Levey M, Timm S, Metler-Altmann T, Luca Borghi G, Koczor M, Arrivault S, Pm Weber A, Bauwe H, Gowik U, Westhoff P. (2019). Efficient 2-phosphoglycolate degradation is required to maintain carbon assimilation and allocation in the C4 plant Flaveria bidentis. Journal of Experimental Botany 70:575–587

Mackinder LCM, Chen C, Leib RD, Patena W, Blum SR, Rodman M, Ramundo S, Adams CM, Jonikas MC (2017) A spatial interactome reveals the protein organization of the algal CO_2_-concentrating mechanism. Cell 171: 133–147 e14

Markelova AG, Sinetova MP, Kupriyanova EV, Pronina NA (2009) Distribution and functional role of carbonic anhydrase Cah3 associated with thylakoid membranes in the chloroplast and pyrenoid of *Chlamydomonas reinhardtii*. Russ J Plant Physiol 56: 761–768

Martinez-Garcia M, van der Maarel MJEC. (2016) Floridoside production by the red microalga *Galdieria sulphuraria* under different conditions of growth and osmotic stress. AMB Express. 6(1):71. doi: 10.1186/s13568-016-0244-6

Moroney JV, Bartlett SG, Samuelsson G. (2001) Carbonic anhydrases in plants and algae. Plant, Cell and Environment 2:141–153

Moulin P, Andría JR, Axelsson L, Mercado JM. (2011) Different mechanisms of inorganic carbon acquisition in red macroalgae (Rhodophyta) revealed by the use of TRIS buffer. Aquatic Botany 95(1):31–38

Nikolau BJ, Ohlrogge JB, Wurtele ES (2003) Plant biotin-containing carboxylases. Archives of Biochemistry and Biophysics 414: 211–222

Noda H. (1993) Health benefits and nutritional properties of nori. J Appl. Phycol. 5:255–258

Patron NJ, Keeling PJ. (2005) Common evolutionary origin of starch biosynthetic enzymes in green and red algae. Journal of Phycology 41 (6): 1131–1141

Raven JA (1995) Photosynthetic and non-photosynthetic roles of carbonic anhydrase in algae and cyanobacteria. Phycologia 34(2):93–101

Rioux L-E, Turgeon SL (2015) Seaweed carbohydrates. In: Tiwari B, Troy D, eds. Seaweed sustainability: food and non-food applications. Amsterdam: Elsevier, 141–192

Shao Z, Xie X, Liu X, Zheng, Z, Huan L, Zhang B, Wang G. (2022) Overexpression of mitochondrial γCAL1 reveal a unique photoprotection mechanism in intertidal resurrection red algae through decreasing photorespiration. Algal Research 66 102766

Viola R, Nyvall P, Pedersén M (2001) The unique features of starch metabolism in red algae. Proc. R. Soc. Lond. B 268, 1417–1422

Von Caemmerer S, Quinn V, Hancock NC, Price GD, Furbank RT, Ludwig M. (2004) Carbonic anhydrase and C4 photosynthesis: a transgenic analysis. Plant, Cell and Environment 6:697–703

Wang D, Yu X, Xu K, Bi G, Cao M, Zelzion E, Fu C, Sun P, Liu Y, Kong F et al. (2020) *Pyropia yezoensis* genome reveals diverse mechanisms of carbon acquisition in the intertidal environment. Nat Commun 11: 4028

Wang S(2004) The ultrastructure of *Porphyra yezoensis*. In Wang S, Pei L, Duan D. eds. The ultrastructure of common red seaweeds in China. Ningbo Publishing Press, Zhejiang, China. 6–9

Wang Y, Stessman FJ, and Spalding MH. (2015) The CO_2_ concentrating mechanism and photosynthetic carbon assimilation in limiting: how *Chlamydomonas* works against the gradient. Plant J. 82:429–448

Wei L, Xin Y, Wang Q, Yang J, Hu H, Xu J (2017) RNAi-based targeted gene knockdown in the model oleaginous microalgae *Nannochloropsis oceanica*. Plant J. 89: 1236–1250

Wei L, El Hajjami M, Shen C, You W, Lu Y, Li J, Jing X, Hu Q, Zhou W, Poetsch A, et al. (2019) Transcriptomic and proteomic responses to very low CO_2_ suggest multiple carbon concentrating mechanisms in *Nannochloropsis oceanica*. Biotechnol Biofuels 12: 168

Wu S, Huang A, Zhang B, Huan L, Zhao P, Lin A, Wang G. (2015) Enzyme activity highlights the importance of the oxidative pentose phosphate pathway in lipid accumulation and growth of *Phaeodactylum tricornutum* under CO_2_ concentration. Biotechnology for Biofuels.8:78, DOI 10.1186/s13068-015-0262-7

Yamano T, Toyokawa C, Shimamura D, Matsuoka T, Fukuzawa H (2021) CO_2_-dependent migration and relocation of LCIB, a pyrenoid-peripheral protein in *chlamydomonas reinhardtii*. Plant Physiol 188: 1081–1094

Young JN, Rickaby REM, Kapralov MV. Filatov DA. (2012). Adaptive signals in algal Rubisco reveal a history of ancient atmospheric carbon dioxide. Philos. Trans. R. Soc. Lond. B Biol. Sci. 367, 483–492

Yu Y, Jia X, Wang W, Jin Y, Liu W, Wang D, Mao Y, Xie C, Liu T. (2021) Floridean starch and floridoside metabolic pathways of *Neoporphyra haitanensis* and their regulatory mechanism under continuous darkness. Marine Drugs 19 (12): 1–19.

Zaffagnini M, Fermani S, Costa A, Lemaire SD, Trost P. (2013) Plant cytoplasmic GAPDH: redox post-translational modifications and moonlighting properties. Front Plant Sci. 4:450. doi: 10.3389/fpls.2013.00450.

Zeebe RE. (2011) On the molecular diffusion coefficients of dissolved CO_2_, HCO_3_−, and CO32− and their dependence on isotopic mass. Geochimica et Cosmochimica Acta 75: 2483–2498.

Zhang B, Xie X, Liu X, He L, Sun Y, Wang G (2020) The carbonate concentration mechanism of *Pyropia yezoensis* (Rhodophyta): evidence from transcriptomics and biochemical data. BMC Plant Biol. 20: 424. https://doi.org/10.1186/s12870-020-02629-4

Zhang B, Liu X., Huan L, Shao Z, Zheng Z, Wang G (2022) Carbonic anhydrase isoforms of *Neopyropia yezoensis*: intracellular localization and expression profiles in response to inorganic carbon concentration and life stage. Journal of Phycology. 58:657–668. https://doi.org/10.1111/jpy.13276

Zheng Z, He B, Xie X, Wang G (2021) Co-suppression in *Pyropia yezoensis* (Rhodophyta) Reveals the Role of PyLHCI in Light Harvesting and Generation Switch. J Phycol 57: 160–171

Zhou W, He L, Yang F, Lin A, Zhang B, Niu J, Wang G (2014) *Pyropia yezoensis* can utilize CO_2_ in the air during moderate dehydration. Chin J Ocean Limnol 32: 358–364

Zhurikova EM., Ignatova LK, Rudenko NN, Mudrik VA, Vetoshkina DV, Ivanov BN (2016) The participation of two carbonic anhydrases of alpha family photosynthetic reactions in *Arabidopsis thaliana*. Biochem. Mosc., 81, 1182–1187.

Zou D, Gao K (2002) Photosynthetic bicarbonate utilization in *Porphyra haitanensis* (Bangiales, Rhodophyta). Chinese Science Bulletin 47:1629–1633

